# Spinal Cord Injury Reduces Serum Levels of Fibroblast Growth Factor-21 and Impairs its Signaling Pathways in Liver and Adipose Tissue in Mice

**DOI:** 10.1101/2021.03.04.433932

**Authors:** Xin-Hua Liu, Zachary A. Graham, Lauren Harlow, Jiangping Pan, Daniella Azulai, William A. Bauman, Joshua Yarrow, Christopher P. Cardozo

**Affiliations:** National Center for the Medical Consequences of Spinal Cord Injury, James J. Peters VA Medical Center, Bronx, NY, USA; Department of Medicine, Icahn School of Medicine at Mount Sinai, New York, NY, USA; Department of Rehabilitation and Human Performance, Icahn School of Medicine at Mount Sinai, New York, NY, USA; Research Service, Birmingham VA Medical Center, Malcolm Randall VA Medical Center, North Florida/South Georgia Veterans Health System, Gainesville, FL, USA; Department of Cell, Developmental and Integrative Biology, University of Alabama-Birmingham, Birmingham, AL, Malcolm Randall VA Medical Center, North Florida/South Georgia Veterans Health System, Gainesville, FL, USA; Research Service and Brain Rehabilitation Research Center, Malcolm Randall VA Medical Center, North Florida/South Georgia Veterans Health System, Gainesville, FL, USA; Division of Endocrinology, Diabetes, and Metabolism, University of Florida College of Medicine, Gainesville, FL, USA

**Keywords:** adiponectin, adipose tissue, diabetes, fibroblast growth factor 21, high fat diet, liver, metabolism, spinal cord injury

## Abstract

Spinal cord injury (SCI) results in dysregulation of carbohydrate and lipid metabolism; the underlying cellular and physiological mechanisms remain unclear. Fibroblast growth factor 21 (FGF21) is a circulating protein primarily secreted by the liver that lowers blood glucose levels, corrects abnormal lipid profiles, and mitigates non-alcoholic fatty liver disease. FGF21 acts via activating FGF receptor 1 and ß-klotho in adipose tissue and stimulating release of adiponectin from adipose tissue which in turn signals in the liver and skeletal muscle. We examined FGF21/adiponectin signaling after spinal cord transection in mice fed a high fat diet (HFD) or a standard mouse chow. Tissues were collected at 84 days after spinal cord transection or a sham SCI surgery. SCI reduced serum FGF21 levels and hepatic FGF21 expression, as well as β-klotho and FGF receptor-1 (FGFR1) mRNA expression in adipose tissue. SCI also reduced serum levels and adipose tissue mRNA expression of adiponectin and leptin, two major adipokines. In addition, SCI suppressed hepatic type 2 adiponectin receptor (AdipoR2) mRNA expression and PPARα activation in the liver. Post-SCI mice fed a HFD had further suppression of serum FGF21 levels and hepatic FGF21 expression. Elevated serum free fatty acid (FFA) levels after HFD feeding were observed in post-SCI mice but not in sham-mice, suggesting defective FFA uptake after SCI. Moreover, after SCI several genes that are implicated in insulin’s action had reduced expression in tissues of interest. These findings suggest that downregulated FGF21/adiponectin signaling and impaired responsiveness of adipose tissues to FGF21 may, at least in part, contribute to the overall picture of metabolic dysfunction after SCI.

## Introduction

Spinal cord injury (SCI) causes partial or total interruption of neural signal transmission between the brain and the periphery, thereby limiting physical activity. Chronic SCI, due to an extremely sedentary lifestyle, invariably results in adverse body composition changes characterized by decreased lean mass and increased adiposity. Such changes are associated with metabolic dysfunction (1–5) leading to an increased risk for cardiovascular disease, liver disorders, and type 2 diabetes (T2DM) compared to the general population (6–9). However, the underlying mechanisms for the predisposition of persons with SCI to metabolic perturbations and the associated cardiac and hepatic disorders remain unclear. Several lines of evidence suggest that alterations in the liver or visceral adipose tissue including neutrophil infiltration, macrophage activation, increased expression of pro-inflammatory cytokines and hepatic lipid accumulation (10–13) which often is associated with development of non-alcoholic fatty liver disease (NAFLD) and non-alcoholic steatohepatitis (NASH) (14, 15). Furthermore, persons with SCI have decreased anabolic hormone levels (16) which are associated with abnormal lipid and metabolic profiles. A clear understanding of hepatic pathology that develops after SCI may provide new insight into the development of impairments of fat and carbohydrate metabolism. However, limited studies have addressed the details of pathophysiological events which occur in the liver after SCI. The mechanism underlying dyslipidemia and whether this change precedes, or is the cause of, systemic metabolic dysfunction after SCI is not known. In addition, the potential mediators, if any, that regulate the interaction between hepatic function and other tissues to achieve systemic metabolic homeostasis after SCI have yet to be investigated.

Fibroblast growth factor-21 (FGF21) is a metabolically active peptide primarily secreted by the liver in response to various nutritional, physiological and pathological stimuli (17). FGF21 is an integral component of the hormone network that modulates the action of insulin, leptin, glucagon-like peptide (GLP1), adiponectin, growth hormone, and insulin-like growth factor (18). FGF21 has been shown to regulate carbohydrate and lipid metabolism in the liver, adipose tissue, skeletal muscle, pancreas, and central nervous system (18–20). In animal models, FGF21 corrects multiple abnormalities of metabolism, including those present in persons with insulin resistance and T2DM (21–23). FGF21 reduces levels of serum triglycerides (TG), total cholesterol, and low-density lipoprotein (LDL)-cholesterol but raises the level of high-density lipoprotein (HDL)-cholesterol, all of which changes represent favorable alterations to the lipid profile (19, 21). Transgenic mice overexpressing FGF21 exhibit resistance to the development of high fat diet (HFD)-induced obesity (24). In addition, FGF21 exerts hepato-protective effects by promoting hepatic lipid and free fatty acid (FFA) metabolism, reducing hepatic adiposity, and ameliorating NAFLD and NASH (19, 22, 25–27). Moreover, FGF21 can act on cardiac muscle to protect against heart disease (28) and ameliorates obesity-related inflammation in a rat model reducing inflammatory cytokine production (29). The molecular mechanisms by which FGF21 produces its physiological effects are, however, still being elucidated. FGF21 transcription is induced by peroxisome proliferator-activated receptor (PPAR)-γ and PPAR-α, which are the targets of agents prescribed to treat T2DM (thiazolidinediones) and hyperlipidemia (fibrates), respectively (23, 30).

FGF21 requires the obligate coreceptor β-klotho (KLB) for cellular signaling both *in vivo* and *in vitro* (31). The binding of FGF21 to KLB enables its interaction with the FGF receptor 1 (FGFR1), thus activating intracellular signaling pathways (32). FGF21 signaling is tissue-specific and is chiefly determined by tissue distribution of KLB. Adipose tissue has high levels of KLB expression and is a primary site of FGF21 action (32, 33). In obese individuals, serum FGF21 levels positively correlate with the subcutaneous adipose tissue (SAT) area (34). FGF21 knockout mice show less SAT and are more insulin-resistant when fed a HFD (24, 34). Mechanistically, FGF21 improves systemic insulin sensitivity by promoting the healthy expansion of SAT (34), thereby increasing serum levels of adiponectin, a major adipokine produced by adipose tissue, which mediates the beneficial effects of FGF21 on the liver and muscle (35, 36). Adiponectin signaling is mediated by activation of two types of receptor: type 1 (AdipoR1) is predominantly expressed in skeletal muscle and activates AMP-kinase; the type 2 adiponectin receptor (AdipoR2) is predominantly expressed in the liver and activates the PPARα signaling pathway (37). Of note, the magnitude of FGF21-induced weight loss is reduced in leptin-deficient mice (33), implying a possible requirement of a functional leptin axis for FGF21 action on body weight regulation (38). While the beneficial effects of FGF21 on metabolic homeostasis have been extensively studied in able-bodied individuals, FGF21 expression and function after SCI has not been investigated.

Persons with SCI have been reported to have reduced energy expenditure resulting from a marked reduction in lean tissue mass and in the ability to ambulate and/or exercise (39, 40). On the other hand, recent reports indicated increased total food intake in persons with SCI when compared to individuals who were immobilized for reasons other than SCI (41, 42). In addition, analysis of the pattern of food intake of overweight or obese persons with SCI suggests that there is an increased proportion of total calories consumed from fat (43). These post-SCI dietary changes almost certainly accelerate the accumulation of fat in various body compartments. However, studies that investigated nutritional status in chronic SCI are limited, and knowledge with respect to how macronutrients impact metabolic gene expression and dysregulation of metabolic homeostasis after SCI is lacking. Furthermore, the relationship between serum FGF21 levels and SCI is unknown.

Using a spinal cord transection model, a high fat diet was recently reported to induce glucose intolerance without increasing fat mass or liver weights (44). To understand the possible role of FGF21 in altered fat, carbohydrate and energy metabolism following SCI, blood and tissue samples collected from SCI mice fed either a high fat diet (HFD) or control diet (ConD) were used to determine circulating FGF21 levels and expression of FGF21 mRNA. The cause of the potential perturbations in FGF21 levels and, if altered, the probable impact on lipid and carbohydrate metabolism was investigated by analyses of relevant signaling and molecular changes in fat, liver and muscle.

## Methods

### Animal studies

Procedures with experimental animals were approved by the Institutional Animal Care and Use Committee at the James J. Peters VA Medical Center and were conducted in accordance with all applicable Federal standards for humane animal care and use of laboratory animals. A detailed description of the experimental protocol has been previously published (44). Briefly, three-month-old male C57BL/6 mice were purchased from Charles River. Animals were randomly assigned to 4 groups: two sham-laminectomy (Sham) groups in which the spinal cord was exposed but not transected, and two thoracic spinal cord transection (level T9) groups. At day 84 after surgery, the animals were euthanized. The liver, adipose tissue including inguinal fat, (iFAT) and omental fat (oFAT); and gastrocnemius muscles were carefully removed and immediately frozen in liquid nitrogen. Blood was collected by ventricular puncture, and serum was frozen for subsequent analysis.

### High-fat diet (HFD)

A lard-based HFD was purchased from Research Diets [D12492; 60% fat, 20% protein, 20% carbohydrate, 5.21 kcal/g energy density]. A source and micro-nutrient control chow with a macronutrient content similar to standard rodent chow [D12450J; 10% fat, 20% protein, 70% carbohydrate, 3.82 kcal/g energy density] was fed to the control-diet (ConD) group. Animals were fed standard rodent chow pre-surgery with mice having ad libitum access to HFD and control chows beginning immediately post-surgery. Chow was replaced every 3d.

### Reverse transcriptase quantitative real-time PCR (RT-qPCR)

Total RNA was extracted using RNasey mini kits (QIAGEN, Germantown, MD). Relative expression of mRNA was determined by *q*Rt-PCR, as described previously (45) using a model ViiA7 thermocycler (Applied Biosystems, Foster City, CA). Specific primers and *Tag*Man Universal Master Mixer were purchased from Applied Biosystems. *Tag*Man Reaction Mixes contained 3 μl cDNA (10 ng/μl). For each sample, the determinations were performed in triplicate. Relative mRNA levels were expressed as fold-change using the 2^−ΔΔCt^ method. Data were normalized relative to 18s RNA.

### Preparation of tissue lysates and Western Blotting (WB)

The tissues were homogenized in 500 μl lysate buffer [150 mM sodium chloride, 3.2 mM Na_2_PO_4_, 0.8 mM K_2_PO_4_ (pH 7.4), 1% NP-40, 0.5% sodium deoxycholate, 0.5% sodium dodecylsulfate] using a tissue grinder bead homogenizer (MP Biomedicals Inc., Solon, OH) followed by centrifugation at 14,000 rpm for 5 min. Western blot analysis was performed as described previously (45). Antibodies against phospho-ACC and total ACC were purchased from Cell Signaling Technology (Beverly, MA). Anti-phospho-PPARα, total PPARα and β-tubulin antibodies were obtained from AbCam (Cambridge, MA). Densitometry was determined using Image Lab software (Bio-Rad, Hercules, CA). β-tubulin was used as a loading control. All assays were performed in triplicate. Mean Ct values for the replicates were used in all subsequent calculations.

### Preparation of nuclear extracts and PPARα DNA-binding activity assay

Nuclear extracts were prepared from mouse liver using a commercial kit (Pierce; Rockford, IL). The binding capacity of PPAR*α* to DNA was assayed using a non-radioactive assay kit from AbCam (Cambridge, MA), which uses a sensitive method for detecting specific transcription factor DNA binding activity in the nuclear extracts. The assays were performed according to the manufacturer’s instructions in duplicate with mean values for the two determinations used in all subsequent calculations.

### *Serum preparation and enzyme-linked immunosorbent assay* (ELISA)

The blood samples were collected on ice and allowed to clot before being centrifugation for 20 min at 1000 *g* with the serum separated from the formed blood cell elements. The FGF21, adiponectin and leptin assays were performed using a commercial ELISA kit (R&D System; Minneapolis, MN); The high molecular weight (HMW) adiponectin assay was performed by a commercial kit (Creative Diagnostics; Shirley, NY). Alanine aminotransferase (ALT) and free fatty acid (FFA) assay kits were purchased from MyBioSource Inc. (Santa Ana, CA) and AbCam respectively. All assays were performed according to manufacturer’s instructions. Assays were performed in duplicate with mean values for the two determinations used in all subsequent calculations.

### Glycogen content assay

Tissue glycogen levels were measured using a colorimetric kit (AbCam). Briefly, mouse liver and skeletal muscle tissues were homogenized with 200 μl distilled water on ice. The homogenates were boiled for 10 min followed by centrifugation at 18,000 rpm for 10 min. Supernatant was collected and subjected to assay according to the manufacturer’s instruction. Samples were assayed in duplicate and mean values for the duplicates were used in subsequent calculations.

### Statistics

Data are expressed as mean values ± SEM. The significance of differences between sham and SCI groups, with or without HFD, was determined using Two-Way ANOVA with a Bonferoni test post-hoc. Statistical calculations were performed using Prism software (GraphPad, San Diego, CA). *p* < 0.05 was considered statistically significant.

## Results

### SCI reduced serum FGF21 levels and inhibited FGF21 receptor expression in adipose tissue

To determine whether FGF21 plays a role in SCI-induced dysregulation of metabolism, serum FGF21 levels were examined. There was a significant reduction in serum FGF21 levels after 84 days of SCI in mice fed a ConD (SCI-ConD) compared to sham-mice fed the same diet (sham-ConD) (Fig. 1A). Because circulating FGF21 levels correlate well with hepatic gene expression (46), FGF21 mRNA expression in mouse liver was determined. As shown in Fig. 1B, SCI led to a significant downregulation of hepatic FGF21 mRNA expression. In addition, we observed that sham mice fed with HFD (sham-HFD) also had reduced FGF21 serum levels and hepatic FGF21 mRNA expression; serum FGF21 and hepatic FGF21 mRNA were further decreased in SCI-HFD mice compared to SCI-ConD mice (Fig. 1B). Adipose tissue expresses high levels of FGFR1 and KLB and is a primary site of FGF21 action in regulating fat and carbohydrate metabolism (32). Therefore, the effects of SCI with or without HFD on FGFR1 and KLB gene expression were examined in adipose tissues. Because increased intra-abdominal fat worsens insulin resistance, as opposed to extra-abdominal fat depots (47), both inguinal fat (iFAT) and omental fat (oFAT) were analyzed. There were significant reductions in both FGFR1 and KLB mRNA expression in iFAT (Fig. 1C, D) and oFAT of SCI-ConD mice compared to sham-ConD mice. While HFD-feeding did not change FGFR1 and KLB mRNA levels in iFAT of sham mice, a significant increase in mRNA expression was seen in oFAT of sham-HFD mice (Fig. 1C, D). In SCI-HFD mice, KLB mRNA levels were reduced compared to SCI-ConD mice in iFAT (Fig. 1C) but not oFAT (Fig. 1D).

**Figure 1.**
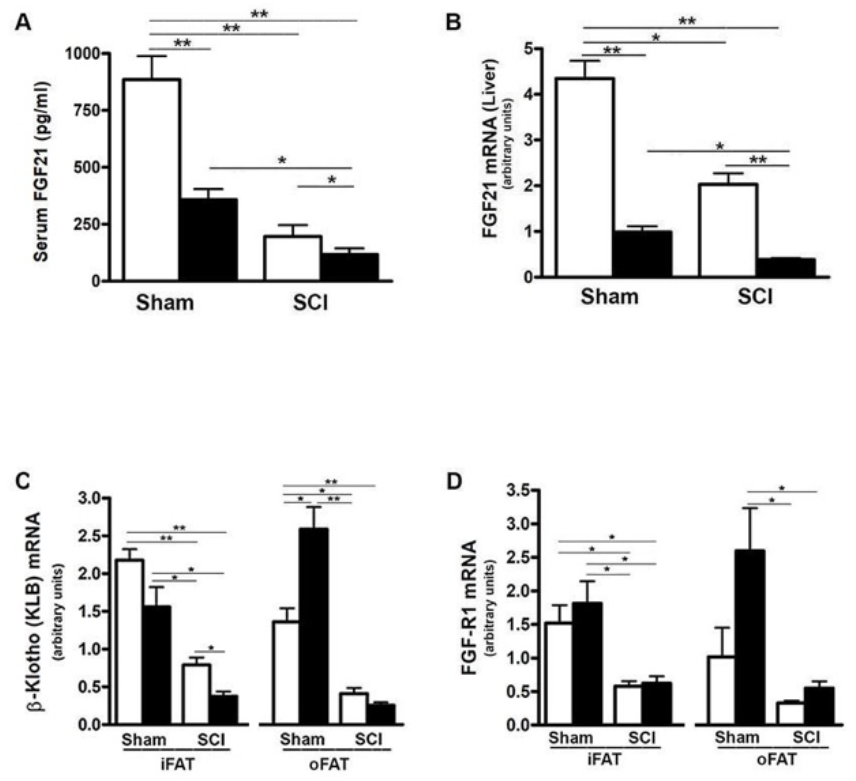
SCI and high-fat diet (HFD) downregulated FGF21 signaling. Mice were fed with either ConD (White Bar) or HFD (Black Bar) after surgery. *(A),* mouse FGF21 serum levels measured by ELISA. *(B),* hepatic FGF21 mRNA expression by Rt-PCR. *(C),* KLB mRNA expression in mouse adipose tissues by Rt-PCR, and *(D),* FGFR1 mRNA expression in mouse adipose tissues by Rt-PCR. Data shown are mean values ± SEM (n = 5). * *p* < 0.05; ** *p* < 0.01.

### SCI reduced expression and serum levels of adiponectin and leptin

It has been demonstrated that FGF21 increases insulin sensitivity through direct effects on fat to stimulate expression and secretion of adiponectin, a critical mediator of FGF21 actions in the liver and skeletal muscle which are two major sites of insulin-induced glucose uptake, and also by specific expansion of subcutaneous fat (34, 35, 36). Thus, we determined whether reduced serum FGF21 impacted the production of adiponectin and leptin as well as adipose tissue differentiation markers. Consistent with reduced levels of FGFR1 and KLB noted in both iFAT and oFAT of SCI-ConD mice, there was a significant decrease in mRNA expression of adiponectin in iFAT and oFAT of SCI-ConD mice as compared to sham-ConD mice (Fig. 2A). Serum levels of both total adiponectin and HMW adiponectin, the most potent form of the protein (48), were also reduced by SCI (Fig. 2B & 2C). Levels of serum total adiponectin in sham-HFD and SCI-HFD mice were further decreased as compared to sham-ConD and SCI-ConD groups, respectively (Fig. 2B). HMW adiponectin was decreased in sham-HFD mice when compared to sham-ConD mice but was not different when comparing SCI-HFD and SCI-ConD mice (Fig. 2C).

**Figure 2.**
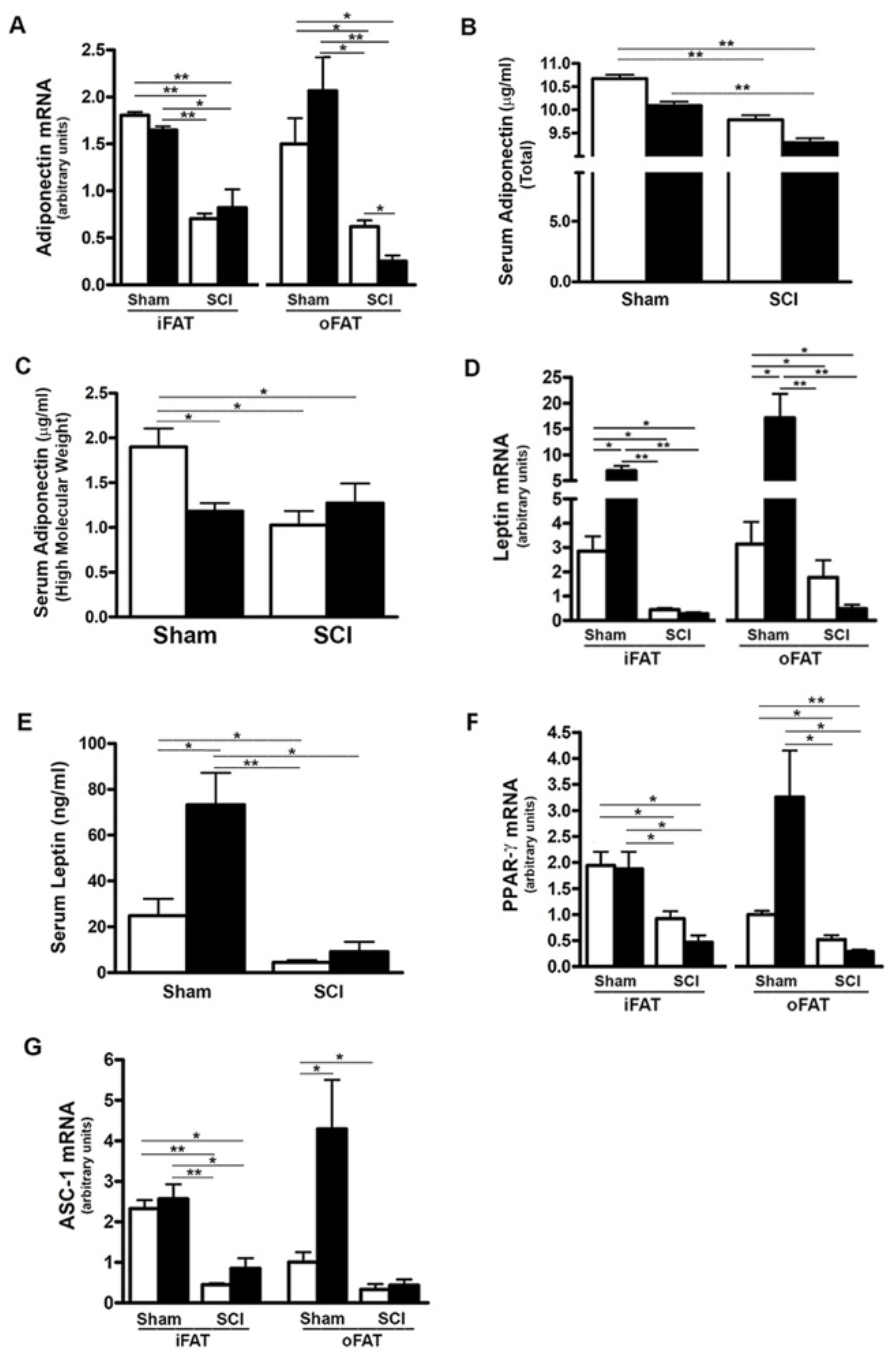
Effects of SCI and HFD on Gene Expression of Adipokines and Differential Markers in Mouse Adipose Tissue. Mice were fed with either ConD (White Bar) or HFD (Black Bar) after surgery. Total RNA was isolated from mouse adipose tissues and subjected to Rt-PCR analysis for *(A),* adiponectin mRNA expression; *(B),* serum total adiponectin levels measured by ELISA; *(C),* serum HMW adiponectin levels measured by ELISA; *(D),* leptin mRNA expression; *(E),* serum leptin secretion measured by ELISA; *(F),* PPARγ mRNA expression; and *(G),* ASC-1 mRNA expression. Data shown are mean values ± SEM (n = 5). * *p* < 0.05; ** *p* < 0.01.

Leptin, another major adipokine, has been implicated in one of the actions of FGF21 (33, 38). The effects of SCI and HFD on leptin mRNA expression and protein secretion were therefore examined. Leptin mRNA in both iFAT and oFAT (Fig. 2D), and serum protein levels (Fig. 2E) were reduced in SCI-ConD mice as compared to sham-ConD mice. Feeding mice with a HFD induced a significant increase in iFAT and oFAT leptin mRNA and serum leptin protein in sham mice, but did not alter serum leptin or leptin mRNA in iFAT or oFAT of SCI-HFD mice when compared to SCI-ConD mice (Fig. 2D, E), suggesting to an impaired ability of adipose tissue to secrete leptin in response to a HFD.

PPARγ a downstream target of FGF21 in adipocytes, is a master transcriptional regulator of adipocyte differentiation, and a determinant of adiponectin expression (23, 49). There was a significant reduction in PPARγ mRNA expression in iFAT and oFAT of SCI-ConD mice as compared to sham-ConD mice (Fig. 2F). Dramatically reduced mRNA levels of ASC-1, a marker of white adipose tissue (50), was also observed in iFAT and oFAT of SCI-ConD mice (Fig. 2G). Expression of PPARγ and ASC were not significantly changed by HFD in sham or SCI mice (Fig. 2F, G). These data indicate that SCI decreased secretion of adiponectin and leptin, and inhibited adipocyte differentiation suggesting a linkage of these impairments to reduced FGF21 signaling.

### SCI reduced hepatic adiponectin receptor 2 (AdipoR2) expression and PPARα activity

AdipoR2 is predominantly expressed in the liver and plays an important role in regulating hepatic lipid metabolism (37, 51). To evaluate whether SCI impacts hepatic adiponectin signaling, AdipoR2 expression was examined in the liver of SCI mice. There was a significant reduction in AdipoR2 mRNA expression in SCI-ConD mice compared to sham-ConD mice and these levels were further decreased in SCI-HFD mice as compared to SCI-ConD mice (Fig. 3A). A HFD also decreased AdipoR2 mRNA in livers of sham-HFD mice compared to sham-ConD mice (Fig. 3A).

**Figure 3.**
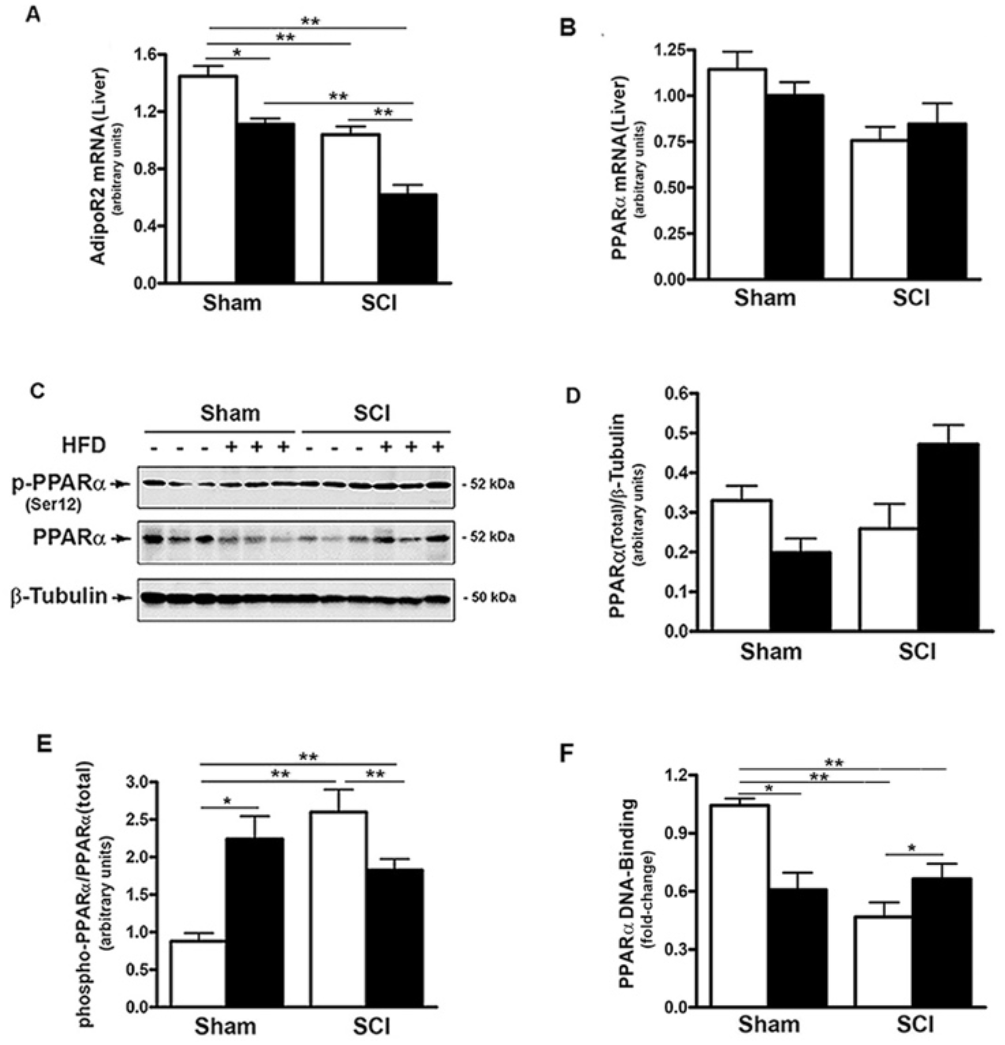
SCI suppressed AdipoR2 expression and PPARα activation. Mice were fed with either ConD (White Bar) or HFD (Black Bar) after surgery. Total RNA was isolated from mouse liver and subjected to Rt-PCR analysis for *(A),* AdipoR2 mRNA; *(B),* PPARα mRNA. *(C),* total protein lysates were extracted from mouse liver and subjected to Western blot analysis using anti-endogenous PPARα or anti-phospho-PPARα (Ser12), or β-tubulin antibodies; (*D),* blots were quantified by scanning densitometry and blots in the middle panel of *C* were normalized relative to β-tubulin (lower panel); *E)*, blots in the upper panel of *C* were normalized relative to endogenous (total) PPARα protein (middle panel). *F),* Nuclear extracts were prepared from mouse liver and subjected to PPARα DNA-binding assay. Data shown are means values ± SEM (n = 5). Data shown in *C* are representative Western blots; * *p* < 0.05; ** *p* < 0.01.

The expression and activity of PPARα, a nuclear receptor that serves as a downstream target of AdipoR2 (37, 52), was also examined. Neither SCI nor HFD changed hepatic PPARα mRNA levels (Fig. 3B). PPARα transcriptional activity is decreased by phosphorylation at Ser12, (53). To understand how SCI altered PPARα activity, the effect of SCI on PPARα phosphorylation and nuclear levels were determined. While SCI had no significant effect on total PPARα protein (Fig. 3C, D). the levels of phospho-(Ser12)-PPARα was increased in SCI-ConD compared to sham-ConD groups (Fig. 3C, E). Phospho-(Ser12)-PPARα was lower in SCI-HFD mice compared to SCI-ConD mice and, by contrast, was higher in sham-HFD mice than sham-ConD mice (Fig. 3C, E). To determine nuclear PPARα activity in SCI vs. sham mice, a DNA-binding assay was performed using nuclear extracts of mouse liver. PPARα DNA-binding capacity was reduced in SCI-ConD mice compared to sham-ConD mice but was elevated in SCI-HFD compared to SCI-ConD mice; in contrast, sham-HFD demonstrated decreased nuclear PPARα compared to sham-ConD (Fig. 3F). These data are consistent with hepatic resistance to adiponectin after SCI due to lower expression of AdipoR2, lower nuclear PPARα and perturbed response of PPARα to dietary fats (52, 54).

### SCI influenced rate limiting steps in hepatic cholesterol and fatty acid metabolism

Because FGF21 regulates hepatic lipid and fatty acid metabolism and reduced PPARα activity after SCI (17, 19, 25–27), the effects of SCI on hydroxy-methyl-glutaryl Co-A reductase (HMGCR), the rate limiting enzyme in cholesterol synthesis, **and ABCA1, the cholesterol efflux regulatory protein, were tested.** HMGCR mRNA levels were markedly increased in livers from SCI-ConD mice compared to sham-ConD mice, whereas reduced mRNA levels were observed in SCI-HFD mice as compared to SCI-ConD mice. HMGCR mRNA levels were also reduced by HFD in sham mice (Fig. 4A). **In contrast, neither SCI nor HFD had no effect on ABCA1 mRNA expression (Fig. 4B) suggesting that SCI may influence hepatic cholesterol synthesis but not efflux.** We next examined the effect of SCI on acetyl-coenzyme A carboxylase (ACC), the rate limiting enzyme controlling fatty acid synthesis and metabolism. Expression of ACCα mRNA levels, the predominant isoform in the liver, was not changed in the SCI-ConD group compared to the sham-ConD group (Fig. 4C). ACCα mRNA was significantly reduced in the sham-HFD and SCI-HFD groups compared to sham-ConD and SCI-ConD groups, respectively (Fig. 4C). Expression of ACCα protein was upregulated in SCI-ConD mice compared to sham-ConD mice without a further change in the SCI-HFD mice when compared to SCI-ConD mice; HFD did not alter total ACCα protein in sham mice (Fig. 4D, E). To understand how SCI and HFD influenced ACCα activity, ACCα phosphorylation at Ser79, which inhibits ACC activity (55), was tested. There was a significant decrease in Ser79-phosphorylated ACCα protein in SCI-ConD mice compared to sham-ConD mice (Fig. 4D, F). Ser79-phosphorylation was not altered by HFD in sham or SCI mice (Fig. 4D, F). These results suggest that SCI suppressed FGF21 signaling associated with increased activity of HMGCR and ACCα pathways that determine hepatic lipid synthesis and metabolism.

**Figure 4.**
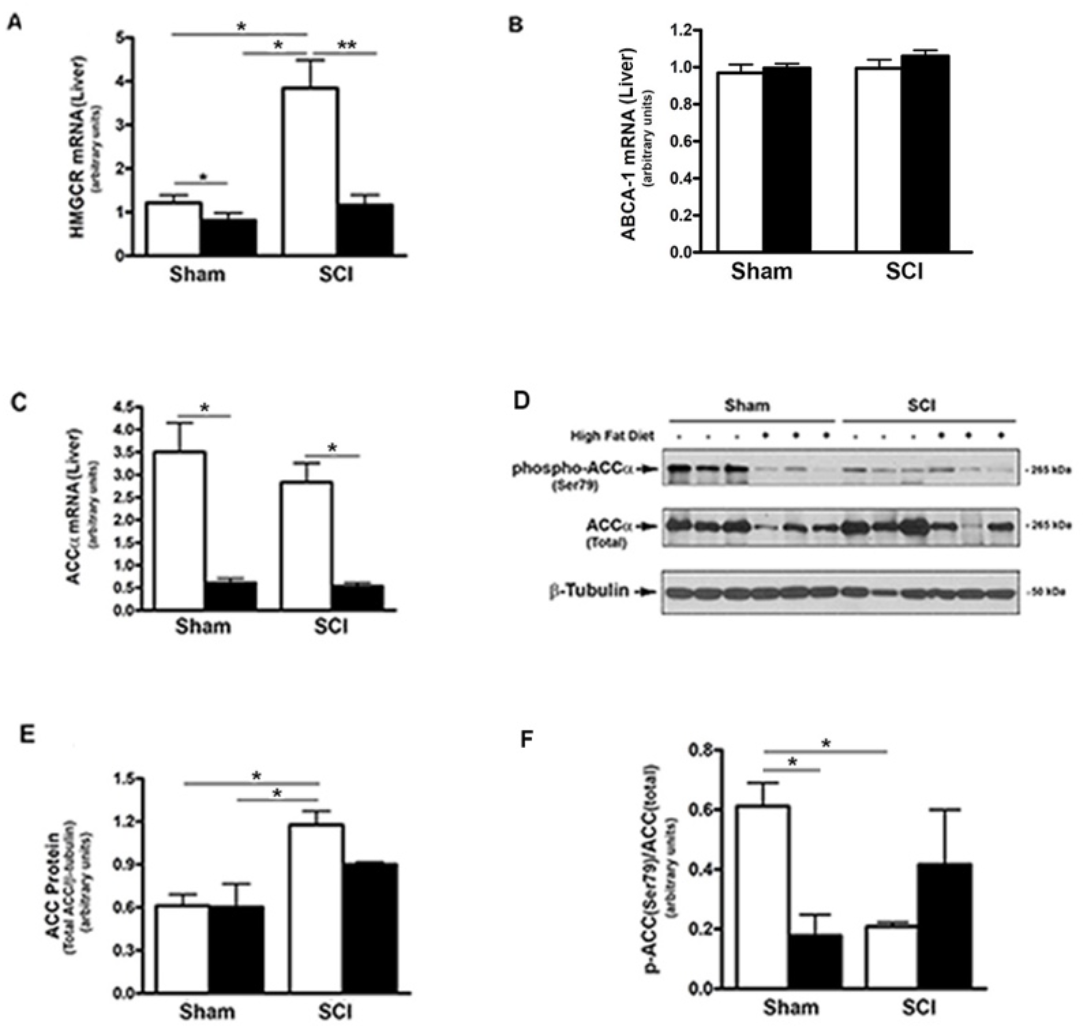
Effects of SCI and HFD on Metabolic Gene Expression in Mouse Liver. Mice were fed with either ConD (White Bar) or HFD (Black Bar) after surgery. Total RNA was isolated from mouse liver and subjected to Rt-PCR analysis for *(A)*, HMGCR mRNA; *(B),* ABCA1 mRNA; *(C),* ACCα mRNA; *(D),* total protein lysates were extracted from mouse liver and subjected to Western blot analysis using anti-ACCα (endogenous) or anti-phospho-ACCα (T172) antibodies; *(E),* blots were quantified by scanning densitometry and blots in the middle panel of *D* were normalized relative to β-tubulin (lower panel); *(F),* blots in *D* (upper panel) were normalized relative to endogenous (total) ACCα protein (middle panel). Data shown are means values ± SEM (n = 5). Data shown in *C* are representative Western blots. * *p* < 0.05, ** *p* < 0.01.

### HFD raised serum FFA and hepatic FABP4 expression after SCI

To further determine the influence of SCI on lipid and fatty acid metabolism, serum FFA levels, which correlate with insulin resistance (56) were examined. When compared to sham-ConD mice, serum FFA levels were not significantly different in SCI-ConD mice although a trend toward lower values in SCI-ConD mice was noted (Fig. 5A). However, as compared to SCI-ConD mice, there was a significant elevation in serum FFA level in SCI-HFD mice, whereas HFD did not alter serum FFA in sham mice (Fig. 5A), suggesting defective FFA uptake by adipose or other tissues after SCI. In addition, the hepatic expression of FABP4 mRNA was examined, the level of which has been linked to insulin resistance, T2DM, hypertension and cardiac dysfunction (57). Although SCI-ConD mice had no change in FABP4 mRNA expression as compared to sham-ConD mice, SCI-HFD mice showed increased expression in hepatic FABP4 mRNA when compared to SCI-ConD mice; HFD did not change FABP4 mRNA levels in livers of sham mice (Fig. 5B). This finding further implies that HFD may worsen SCI-induced metabolic dysfunction.

**Figure 5.**
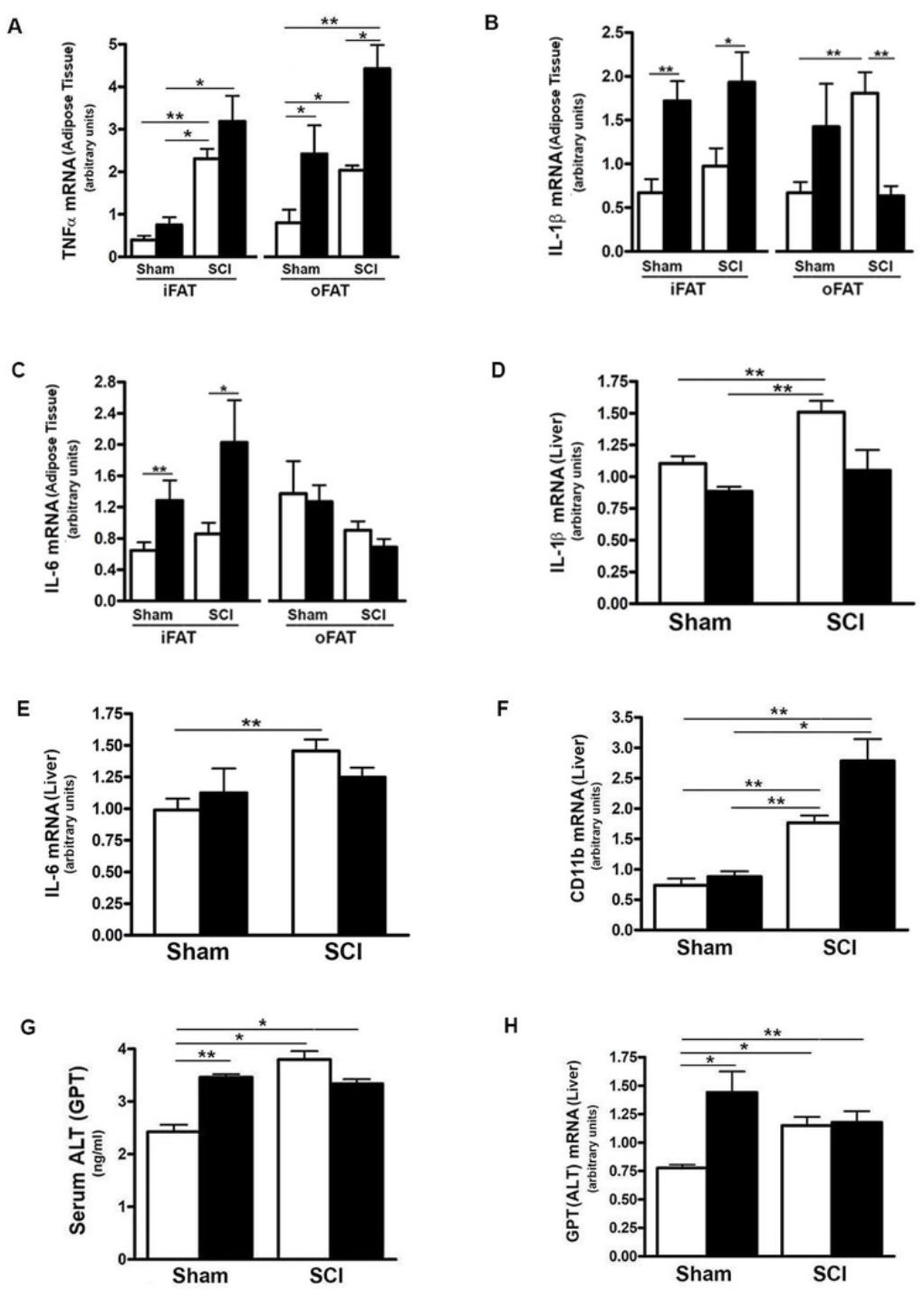
Differential Effect of HFD on Serum FFA Level in Post-SCI and Sham Mice. Mice were fed with either ConD (White Bar) or HFD (Black Bar) after surgery. *(A),* mouse serum was collected and subjected to ELISA for FFA serum level assay; *(B),* total RNA was isolated and subjected to Rt-PCR for hepatic FABP4 mRNA expression. Data shown are means values ± SEM (n = 5). * *p* < 0.05; ** *p* < 0.01.

### SCI induced increased inflammatory gene expression in fat and liver

Inflammatory changes in liver and fat have been linked to impaired metabolism of carbohydrates (58). To understand if SCI and/or HFD influenced inflammatory status in liver or fat, selected mRNA levels of inflammatory cytokines were determined. Expression of TNFα mRNA in iFAT and oFAT was increased in SCI-ConD mice as compared to sham-ConD mice; a further increase in TNFα mRNA levels were observed in SCI-HFD mice when compared to SCI-ConD mice (Fig. 6A). In sham mice, HFD increased TNFα mRNA expression in oFAT but not iFAT (Fig. 6A). The expression levels of IL-1b and IL-6, two major cytokines induced by inflammation, were also tested. As shown in Fig. 6, SCI had no effect on IL-1β or IL-6 mRNA in iFAT, but significantly upregulated IL-1β levels in oFAT (Fig. 6B). HFD-feeding increased IL-1β and IL-6 mRNA levels in iFAT but not in oFAT (Fig. 6B, C). While SCI had no effect on TNFα or TNF-receptor mRNA expression in mouse liver (Supplemental Data, Fig. 2), significant upregulation of expression of IL-1β (Fig. 6D), IL-6 (Fig. 6E) and CD11b mRNA, a macrophage marker, (Fig. 6F) were observed in the liver from SCI-ConD mice as compared to sham-ConD mice. The expression of IL-1β and IL-6 mRNA was not changed by HFD in sham or SCI mice. In contrast, while CD11b mRNA was not changed in sham-HFD mice, it was significantly elevated in SCI-HFD mice as compared to SCI-ConD mice (Fig. 6F).

**Figure 6.**
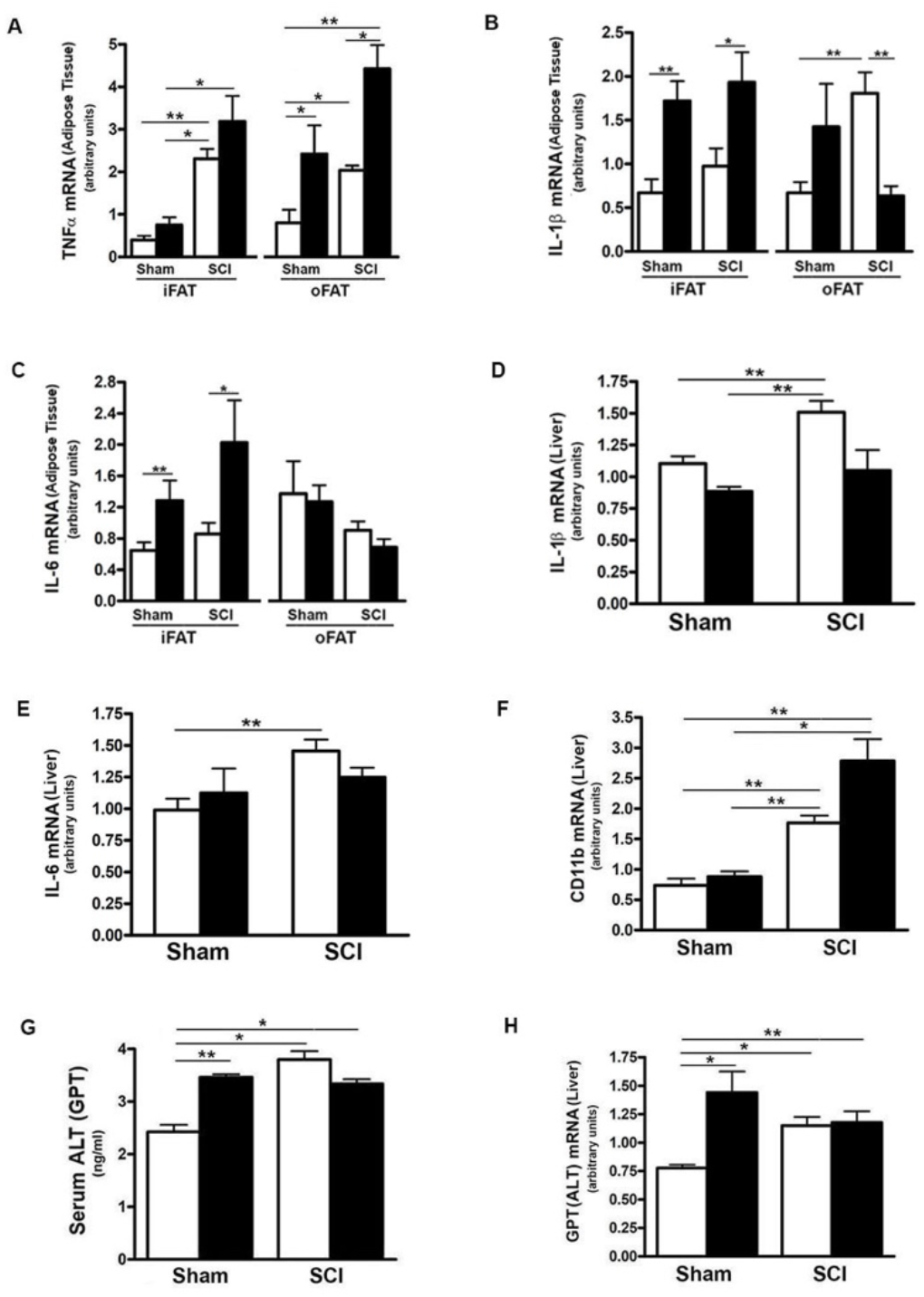
SCI Caused Hepatic Pro-Inflammatory State and Liver Toxicity. Mice were fed with either ConD (White Bar) or HFD (Black Bar) after surgery. *(A),* total RNA was isolated from mouse adipose tissues and subjected to Rt-PCR for TNFα mRNA expression; *(B),* IL-1β mRNA in adipose tissue; *(C)*, IL-6 mRNA expression in adipose tissue; *(D)*, total RNA was isolated from mouse liver and subjected to Rt-PCR for IL-1β mRNA; *(E)*, hepatic IL-6 mRNA; *(F)*, hepatic CD11b mRNA expression; *(G),* Mouse serum was collected and subjected to ELISA for ALT serum level; *(H),* total RNA was isolated from mouse liver and subjected to Rt-PCR for hepatic ALT mRNA expression. Data shown are means values ± SEM (n = 5). * *p* < 0.05; ** *p* < 0.01.

To understand the potential impact of elevated CD11b on hepatocytes, serum and hepatic mRNA levels of ALT, a hepatic enzyme that has been used as a sensitive marker of liver injury, were determined. Elevated levels of both serum ALT (Fig. 6G) and expression of hepatic ALT (Fig. 6H) were observed in SCI-ConD mice as compared to sham-ConD mice. Serum ALT was increased in sham-HFD mice compared to sham-ConD mice but was not different between SCI-HFD and SCI-ConD mice (Fig. 6G). Expression of hepatic ALT mRNA was increased in sham-HFD mice compared to sham-ConD mice, but not changed in SCI-HFD mice when compared to SCI-ConD mice (Fig. 6H). Taken together, these findings suggest that SCI may trigger inflammatory responses marked by increased production of cytokines from adipose tissue and enhanced numbers of macrophages migrating to the liver, which could result in hepatocellular injury and liver dysfunction, as well as contribute to the general metabolic perturbations observed after SCI.

### SCI resulted in tissue specific reduction of insulin signaling-related gene expression

Because there are links among FGF21, adiponectin signaling, hepatic inflammation and insulin action (22, 25, 27, 36, 59), we determined whether the dysregulated FGF21 and adiponectin signaling observed after SCI were associated with altered mRNA expression of selected components of the insulin signaling pathway. As compared to sham-ConD, SCI-ConD mice demonstrated reduced expression of IRS-1 mRNA in iFAT and oFAT (Fig. 7A), as well as in liver (Fig. 7C). When comparing SCI-ConD and sham-ConD mice, decreased Glut4 mRNA expression was seen in iFAT and oFAT (Fig. 7B). It is appreciated that muscle is the primary site of glucose uptake and disposal. In gastrocnemius muscle, expression of insulin receptor mRNA level was reduced in SCI-ConD mice as compared to sham-ConD mice (Fig. 7E). However, there was no significant difference in Glut4 or IRS-1 mRNA in gastrocnemius muscle from SCI-ConD as compared to sham-ConD mice, and these parameters were not altered by HFD (Supplemental Data, Figure 1). To further evaluate how impaired FGF21 and adiponectin pathways impact insulin signaling, glycogen content was measured in mouse liver and skeletal muscle. There was a significant reduction in glycogen content in gastrocnemious muscle in SCI-ConD compared to sham-ConD mice (Fig. 7F); muscle glycogen levels were not altered by HFD in either sham or SCI mice. In contrast, SCI with or without HFD had no effect on glycogen levels in liver (Fig. 7D).

**Figure 7.**
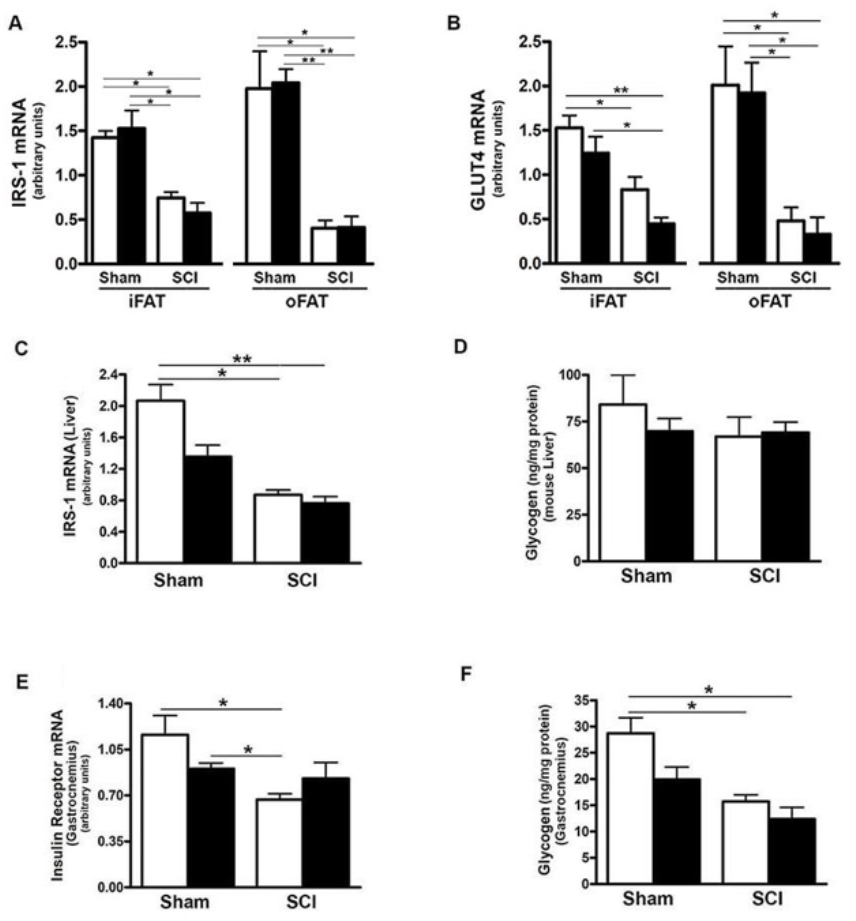
SCI Dysregulated Tissue Specific Expression of Insulin Signaling-Related Genes. Mice were fed with either ConD (White Bar) or HFD (Black Bar) after surgery. Total RNA was isolated from adipose tissues and subjected to Rt-PCR for *(A),* IRS-1 mRNA expression, and *(B),* Glut4 mRNA expression; *(C),* total RNA was isolated from mouse liver and subjected to Rt-PCR for hepatic IRS-1 mRNA expression; *(D),* liver glycogen content measured by colorimetric assay; *(E)*, total RNA was isolated from mouse gastrocnemius and subjected to Rt-PCR for insulin receptor mRNA expression; *(F)*, colorimetric assay for glycogen content in mouse gastrocnemius. Data shown are mean values ± SEM (n = 5) * *p* < 0.05; ** *p* < 0.01.

## Discussion

The central question addressed in this study was whether SCI per se has an impact to change serum FGF21 levels or activity of FGF21-dependent signaling, including that through adiponectin, in a mouse model of complete spinal cord transection. To better understand the meaning of changes in levels of these regulatory factors, the analysis employed a broad approach that examined signaling in liver, iFAT, oFAT and skeletal muscle which, collectively, are responsible for most effects of insulin on blood glucose levels, with adipose tissue and liver also playing key roles in synthesis, storage and release of fats. The principal conclusions of our work herein are that SCI reduced serum FGF21 and adiponectin levels, including total and the more bioactive HMW moiety, and the expression of hepatic FGF21 and adipocyte expression of adiponectin. The data suggest that SCI reduced signaling in adipocytes in response to FGF21 via KLB and FGFR1, and diminished adiponectin action on the liver through reduced hepatic AdipoR2 expression and decreased PPARα activation. In addition, SCI increased levels in liver of inflammatory cytokines, including IL-1β, IL-6 and CD11b, as well as in both iFAT and oFAT of TNFα mRNA, the changes were exacerbated by HFD suggesting that these adipose tissue deposits contribute to the state of low-grade chronic inflammation reported after SCI (58), which was associated with elevated liver expression of ALT mRNA and serum levels of ALT, indicative of hepatocellular injury. Moreover, SCI reduced insulin receptor in gastrocnemius muscle, and reduced mRNA levels for IRS-1 and Glut-4 in iFAT and oFAT, and IRS-1 in liver suggesting impaired insulin signaling pathway after SCI. Evidence of perturbations in regulation of synthesis of steroids, fatty acids and fatty acid metabolism after SCI included greatly elevated HMGCR mRNA expression and ACCα activity in liver after SCI. During administration of a HFD, serum FFA rose in SCI mice on HFD, but not control mice, suggesting impaired ability to store FFA as triglycerides in adipocytes or to use FFA as fuel in muscle and other tissues. FABP4 mRNA was elevated in liver of SCI-HFD mice which is notable because serum levels of the corresponding protein are linked to T2DM and other disorders (52). Taken together, the data indicate defects in metabolism of fats and carbohydrates in multiple tissues after SCI and implicate profound dysregulation of the FGF21 – adiponectin metabolic regulatory circuit as one key driver of these many perturbations.

Hepatic FGF21 expression is regulated by many factors that include nutrients, physical activity and medications (17, 59). Reasons that hepatic FGF21 expression is reduced by SCI are unclear. Because of the key role played by PPARα in activation of FGF21 transcription in liver, the reduced nuclear levels and transcriptional activity of PPARα provide insight with respect to one possible mechanism by which FGF21 mRNA levels are reduced in liver after SCI (60), but other possibilities exist. While it remains uncertain how exercise increases FGF21, the linkage between physical activity and FGF21 suggests that just as exercise raises levels of this hepatokine (61, 62), a sedentary lifestyle may lower them. The possibility should also be considered that cues from hepatic or adipocyte inflammation, which was present in both adipose tissues (iFAT and oFAT) and liver, as suggested by the elevated expression of mRNA levels for inflammatory cytokines, particularly for TNFα in the adipose tissue, and IL-1β, IL-6 and CD11b mRNA in the liver, could lead to reduced hepatic FGF21 expression. These data suggest that an increased focus on derangements in liver function as a key factor in impaired fat and carbohydrate metabolism after SCI should be considered.

The reduction in adipocyte expression of adiponectin mRNA and serum adiponectin level observed in spinal cord transected mice in the current study may be linked in part to the lower level of serum FGF21 observed after SCI; the data also suggest resistance to FGF21 actions based on lower expression of FGFR1 and KLB in iFAT and oFAT cells. While controversial, some evidence for FGF21 resistance has been presented (63). Other explanations cannot be excluded from the data available and include reduced capacity for PPARγ to activate adiponectin transcription as a result of local pro-inflammatory signals (49). Whatever the mechanisms responsible for reduced serum adiponectin after SCI might be, they were coupled with several molecular changes in liver that might be interpreted as indirect evidence of resistance to adiponectin action. These include lower levels of AdipoR2 mRNA and PPARα activity. While not extensively studied, the concept of adiponectin resistance has been proposed previously and some experimental support for its existence has been presented (64).

Lower expression in iFAT and oFAT of leptin and reduced serum leptin levels were also observed. These changes are consistent with a prior report of low serum leptin after a contusion SCI in rats (65). Implications of these changes are unclear but it should be noted that leptin has been implicated in multiple biological responses including of FGF21 action (33, 38). Regulation of leptin expression remains incompletely understood, but it is clear that leptin levels correspond to adipose tissue mass and are reduced by low calorie intake (66). Thus, the low mass of adipose tissue deposits in the SCI mice (44) observed in this investigation may explain, at least in partial, the low leptin levels observed.

Serum FFA levels were significantly elevated in HFD-fed SCI mice compared to SCI-ConD mice but not in sham-HFD mice. These observations suggest impaired ability of SCI mice to clear FFA from the circulation or to metabolize FFA such that they accumulate in blood. Elevated serum FFA levels have been shown to reduce tissue insulin sensitivity through multiple mechanisms that include reduced glucose uptake by skeletal muscle and increased TNFα release by liver Kupfer cells (67, 68), Importantly, elevated FFA levels are a risk factor for insulin resistance that may be independent of total fat mass (56). Elevations in serum FFA in SCI-HFD mice were associated with increased hepatic expression of FABP4. FABP4 is thought to have numerous functions that include its ability to bind free fatty acids and retinoids within cells, to participate in uptake of FFA from the extracellular space, and to route FFA to nuclear PPAR receptors to appropriately regulate the gene expression programs they control (69, 70); the implications of elevated hepatic FABP4 following SCI fed with a HFD are uncertain, but it is noteworthy that elevated serum levels of the protein encoded by this gene showed a positive correlation with high-sensitivity C-reactive protein (a measure of systemic inflammation), lipid parameters, and the homeostatic model assessment of insulin resistance (HOMA-IR) (56, 69–71). Taken together, these findings indicate that following SCI, mice were less tolerant of a HFD, and that this type of diet elicited several significant and deleterious changes.

Effects of SCI on the serum lipid profile have been evaluated in several other studies. Elevated serum triglycerides were observed at one month after SCI in T3-transected rats associated with increased visceral adiposity (72). In a separate study, serum FFA and glucose were elevated 42 days after a 200 kdyne contusion injury in rats (73). While extensive study of serum lipid profiles has been performed in patients with SCI (1), to date these studies have not characterized serum FFA levels.

An unexpected finding was the upregulation in SCI-ConD mice of HMGCR, a rate-limiting enzyme in the synthesis of cholesterol, possibly reflecting increased tissue demand, which might be expected during reparative responses in the spinal cord or to support neuroplasticity. In SCI patients, abnormalities of lipid metabolism include reduced HDL cholesterol (1, 2) and elevated triglycerides in response to a high fat challenge when compared to data for historical controls (74). In rats that were studied 1 month after thoracic level-10 transection, serum triglycerides, total and HDL cholesterol levels tended to be lower as compared to sham-operated controls, although this difference was not significant (72). Further study is needed to understand how SCI impacts metabolism of fats or how SCI perturbs the networks of adipokines and hepatokines that regulate them.

The data presented suggest that at 84 days after spinal transection, there is accumulation of tissue macrophages in the liver, as reflected by increased expression of inflammatory cytokines, including IL-1β, IL-6 and CD11b. There is also some evidence of inflammation in fatty tissue based on elevated expression of TNFα mRNA in both iFAT and oFAT, and increased IL-1β mRNA in oFAT. These alterations would be expected to stress hepatocytes, impair insulin action on hepatocellular regulation of glucose uptake and glucose output and, possibly, injure hepatocytes as well. Evidence of hepatocellular impairment, and possibly injury, was observed in post-SCI mice based on elevated circulating levels of ALT, as well as increased hepatic ALT expression. It is noteworthy that Kuppfer cells, the resident hepatic macrophages, have been linked to NAFLD, a risk factor for insulin resistance (75), thus indirectly implicating the changes in CD11b to liver disease and insulin resistance after SCI. Elevated cytokine expression in fat deposits is also thought to be a risk factor for impaired glucose homeostasis (76, 77), suggesting that such alterations in adipose tissues observed in the current study may further contribute to impaired glucose handling and insulin action in SCI mice. This postulate is indirectly supported by findings of reduced expression of several key genes in insulin signaling in iFAT, oFAT, liver and gastrocnemius muscle. However, the data do not establish causal relationship between TNFα expression in adipose tissue and altered fat or carbohydrate metabolism. Also unclear is the cellular source of cytokines in adipose tissues, but prior studies indicate that infiltration of fat deposits by tissue macrophages is the primary explanation for such changes (76). Evidence of hepatic inflammation and hepatocellular injury that increases with time to 21 days after SCI has been observed in rats following a 200 kdyne contusion SCI. In the same study, elevations in serum ALT were observed in rats with cervical (C5), thoracic (T8) and lumbar (T12) contusion injuries (11). In that study, hepatic mRNA levels for IL-1ß increased at days 3 and 7 post-SCI while TNFα increased at 14 and 21 days, suggesting a time-dependent response. Hepatic lipids, assessed by Oil red O staining, were also increased at 14 days post injury (11). A more recent study from the same group found elevated hepatic CD11b at 6-weeks after a 200 kdyne injury at T8 that was associated with increased mRNA levels for several cytokines, including TNFα and IL-1ß, and increased hepatocellular lipids and iron levels (73). These findings are in agreement with the increased expression of CD11b in liver in the current study. However, no significant alteration in hepatic expression of cytokines was observed in the present study. Reasons for these discrepant findings are unclear but may include differences in time since injury, species and/or responses to contusion versus transection SCI.

It was previously reported that, in mice, spinal cord transection results in marked decreases in weights of iFAT and oFAT and that HFD did not increase weights of these fat deposits in SCI mice (44). Similarly, a reduction in total fat mass in rats after transection at thoracic level-4 has been reported and was associated with a small increase in metabolic rate and food intake (78). There is insufficient data to determine why the observed decreases in fat depots occurred. It has been argued that because body weights of the mice from which tissues were obtained for the current study increased over time after SCI that their net caloric intake exceeded metabolic demands. However, it remains unclear why gains in body weight were not associated with a return toward normal adipose tissue mass; possible explanations include impaired ability of adipocytes to take up and store lipids or, possibly, a loss of the normal replacement of aging adipocytes from the pool of adipocyte progenitors such that adipose tissues become depleted of healthy and fully functional adipocytes. Similarly, the reduced levels of PPARγ and ASC-1 observed in the current study could simply reflect low fat mass. Adipocytes undergo turnover whereby there is constant replenishment of aging or damaged adipocytes ones formed from a committed pool of progenitors (79). Thus, failure of HFD to promote gains in adipose tissue mass may indicate damage to adipocytes and/or inadequate replenishment of adipocytes from pools of progenitors. Of note, it was reported that FGF21 knockout in mice displayed disorders of adipose tissue due to defects in PPARγ signaling and PPARγ-dependent gene expression (34, 80). This report demonstrates that it is plausible that the reduced levels of FGFR1 and PPARγ activity observed in iFAT and oFAT could result in dysfunction of adipose tissue, low adipose tissue mass and impaired adiponectin expression and release.

It is now accepted that fat deposits are highly specialized in their functions depending on their location. oFAT is particularly important when considering the pathogenesis of insulin resistance and diabetes, perhaps because adipokines and free fatty acids liberated by oFAT are sent to the liver by the portal circulation. Unexpectedly then, in the present study, gene expression changes in iFAT and oFAT from SCI mice, and responses in SCI mice of these fat deposits to HFD were similar in magnitude and direction suggesting that physiological adaptations to SCI in mice overcame phenotypic differences related to localization of these fat deposits.

It was previously reported that HFD induced glucose intolerance in spinal cord transected mice at 84 days after injury and that fasting glucose was modestly increased in SCI-ConD mice at this time point (44). When these findings are taken together with the ability of FGF21 to improve insulin action, and low FGF21 observed in SCI in mice, it was of interest to examine insulin signaling pathways in tissues from these SCI mice. Whereas no reduction in IRS-1 or Glut-4 mRNA was observed in skeletal muscle, reduced levels of IRS-1 mRNA were observed in iFAT and oFAT, as well as liver, and these changes were not influenced by administration of a HFD. IRS-1 is critical to coupling insulin receptor and downstream signaling pathways while Glut4 represents the primary mechanism for insulin-induced glucose uptake. The findings suggest that, in the spinal cord transected mouse model at least, loss of insulin responsiveness of liver and fat deposits is an important mechanism leading to insulin resistance and diabetes. However, as discussed above, the data do not exclude a possible deterioration of function of adipose tissue as an explanation for the uniform decrease in insulin signaling pathways genes in adipose tissues. Further studies are necessary to better understand the respective roles of fat, liver and muscle in development of T2DM after SCI.

A limitation to be considered when interpreting our findings or considering their translation is that the approach employed did not directly test for physiological significance of lower FGF21 levels in the SCI mouse model used. Consequently, although it is attractive to posit that the changes observed downstream of serum FGF21 may be attributable to the lower serum FGF21 levels observed after spinal cord transection, this mechanistic linkage is yet to be proven. It should also be noted that whereas persons with SCI usually gain body fat mass, mice with spinal cord transection lost fat mass (44). This difference may limit how well changes in the spinal cord transection mouse model used in the current model reflect what occurs in individuals with SCI. Another limitation of these studies is that they evaluated only male mice; males were selected because male C57B6 mice are more sensitive to high-fat-diet-induced glucose intolerance than females. We expect that the pattern of changes in serum and hepatic FGF21 levels would be the same between males and females after SCI although some gender effects on the magnitude of changes can not be excluded.

In summary, this study, the impact of SCI on expression of and signaling by FGF21 was investigated. Major findings of the study are that at 84 days after a complete spinal cord transection, hepatic FGF21 expression and serum FGF21 protein were reduced, associated with lower serum total and HMW adiponectin and leptin. A HFD raised serum FFA in SCI but not sham mice suggesting impaired ability for tissue to take up or metabolize fats. Prior studies demonstrated glucose intolerance in the SCI-HFD mice model that was used for these studies. Decreased expression for insulin signaling-related genes were observed in adipose tissue and liver but, surprisingly, not skeletal muscle. Taken together, reduced levels of FGF21 and adiponectin provide attractive mechanisms for the links between SCI and insulin resistance and T2DM. Moreover, the findings suggest key roles for perturbed function of adipose tissue and liver in the pathogenesis of T2DM after SCI. Further research will be needed to formally prove a causal relationship between abnormal levels of these proteins and carbohydrate and fat metabolism after SCI.

## Acknowledgements

We are very grateful to Dr. Kenneth Cusi for his very insightful discussions about the project.

## Supplemental Data

**Figure 1.**
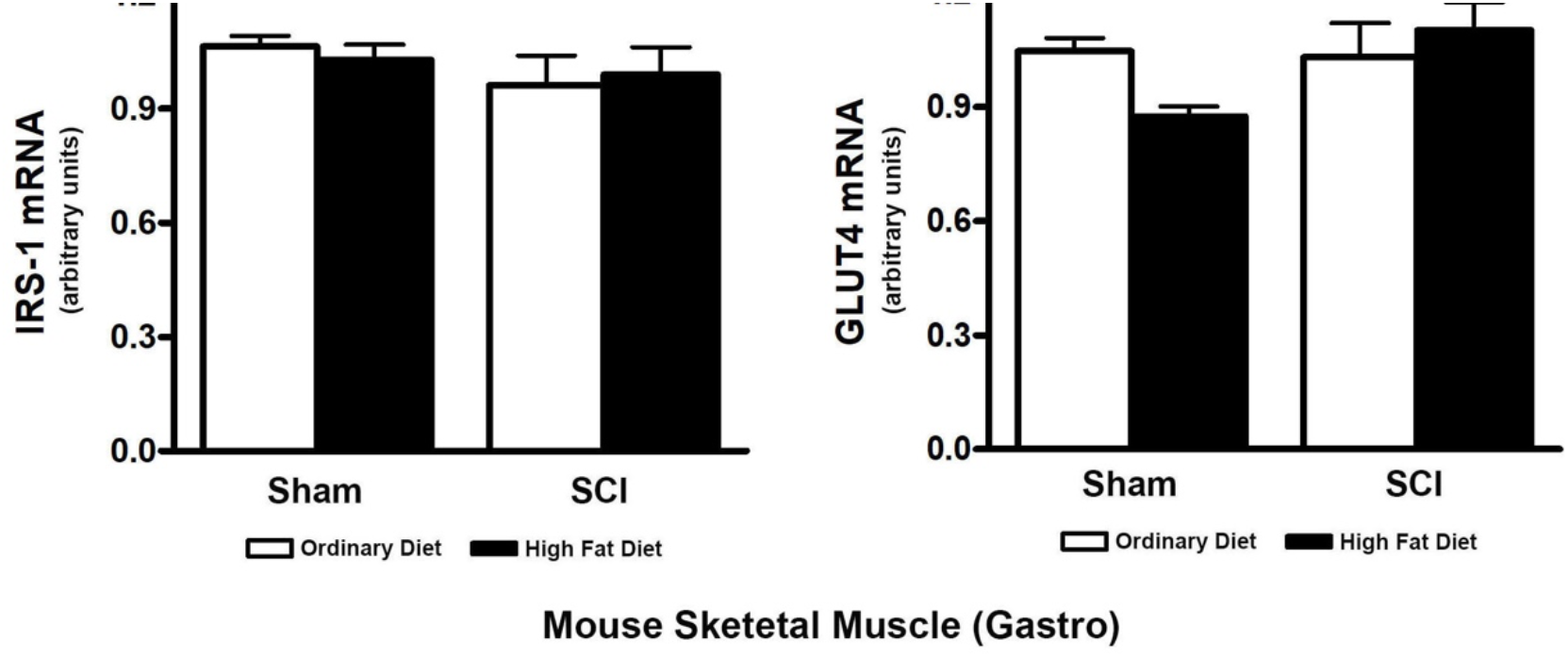
Effect of SCI and HFD on Glut4 and IRS-1 mRNA Expression in Skeletal Muscle. Mice were fed with either ConD (White Bar) or HFD (Black Bar) after surgery. Total RNA was isolated from mouse gastrocnemius and subjected to PCR analysis. *(A),* IRS-1 mRNA expression; and *(B),* Glut4 mRNA expression. Data shown are mean values ± SEM (n = 5).

**Figure 2.**
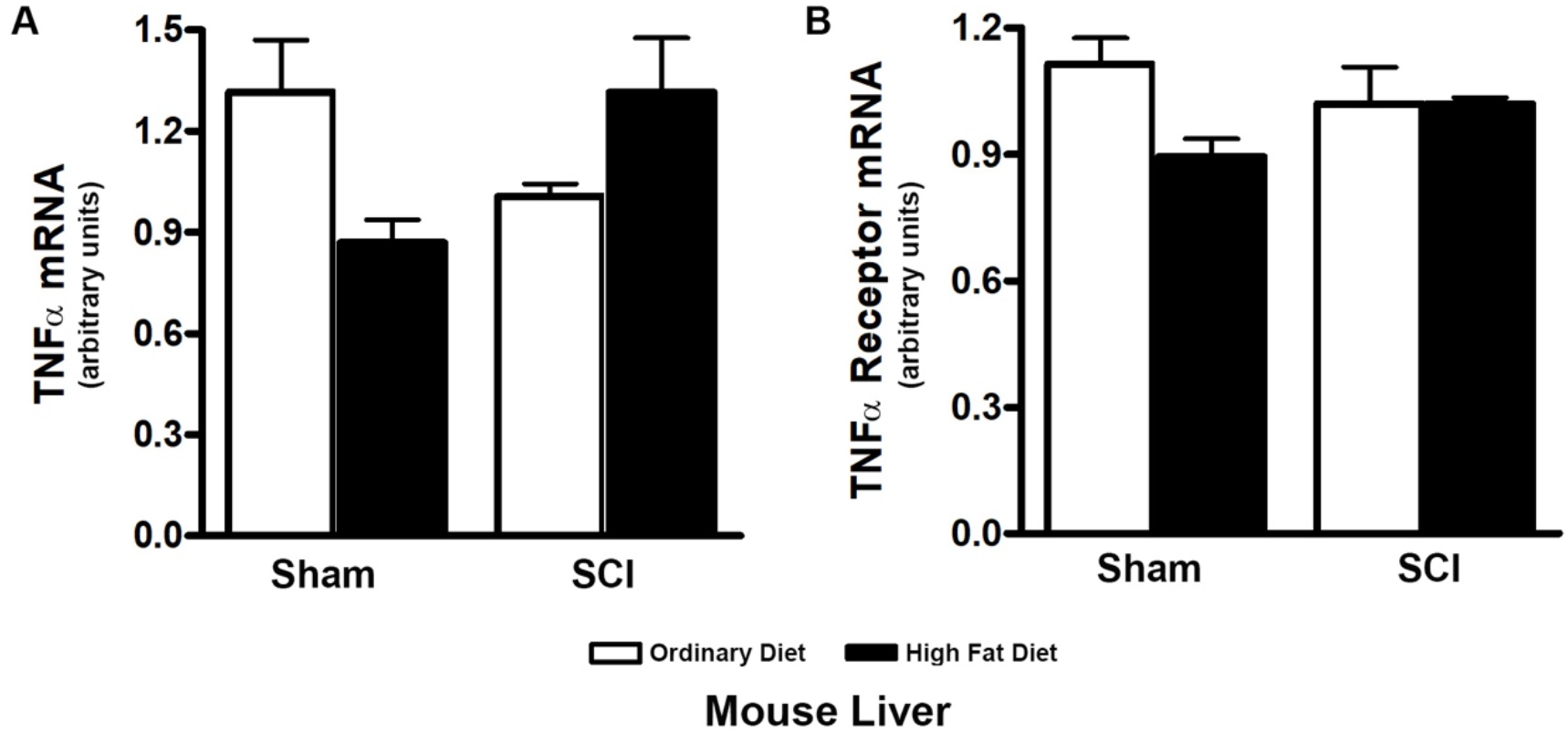
Effect of SCI and HFD on Hepatic TNFα and TNFα receptor mRNA Expression. Mice were fed with either ConD (White Bar) or HFD (Black Bar) after surgery. Total RNA was isolated from mouse liver and subjected to PCR. *(A),* TNFα mRNA expression; and *(B),* TNFα receptor mRNA expression. Data shown are mean values ± SEM (n = 5).

**Figure 3.**
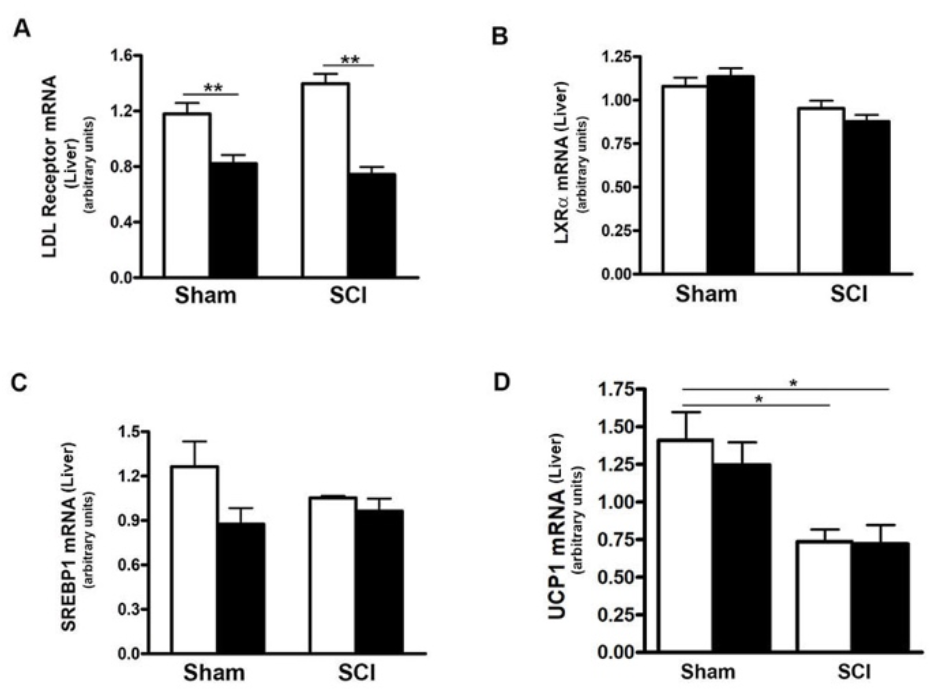
Effects of SCI and HFD on lipid and fatty acid metabolism-related gene expression in the liver. Mice were fed with either ConD (White Bar) or HFD (Black Bar) after surgery. Total RNA was isolated from mouse liver and subjected to PCR. *(A),* LDL receptor mRNA; *(B), Liver X receptor-α* (LXRα mRNA; *(C)*, SREBP1 mRNA; *(D)*, UCP1 mRNA. Data shown are mean values ± SEM (n = 5).

**Table 1.** Summary of Metabolic Gene Expression after SCI with or without HFD

| Gene Name | Tissue Origins | SHAM-ConD | Sham-HFD | SCI-ConD | SCI-HFD |
| --- | --- | --- | --- | --- | --- |
| FGF21 Secreted Protein | Serum | 1.0 | 0.405 ** | 0.222 ** | 0.132 ** # |
| FGF21 mRNA | Liver | 1.0 | 0.219 ** | 0.403 * | 0.084 ** # |
| β-Klotho (KLB) | iFAT | 1.0 | 0.716 * | 0.362 ** | 0.173 ** ## |
| β-Klotho (KLB) | oFAT | 1.0 | 3.071 * | 0.489 * | 0.305 ** |
| FGFR1 | iFAT | 1.0 | 1.195 | 0.382* | 0.411* |
| FGFR1 | oFAT | 1.0 | 2.557 | 0.324 | 0.540 # |
| Adiponectin Protein(Total) | Serum | 1.0 | 0.953 | 0.929 ** | 0.886 ** |
| Adiponectin Protein(HMW) | Serum | 1.0 | 0.628 * | 0.460 * | 0.539 * |
| Adiponectin mRNA | iFAT | 1.0 | 0.914 | 0.390 ** | 0.455 ** |
| Adiponectin mRNA | oFAT | 1.0 | 1.378 | 0.428 * | 0.168 * ## |
| Leptin Secreted Protein | Serum | 1.0 | 2.950 * | 0.178 * | 0.365 |
| Leptin mRNA | iFAT | 1.0 | 2.435 * | 0.156 * | 0.097 * |
| Leptin mRNA | oFAT | 1.0 | 5.450 * | 0.562 | 0.155 * ## |
| PPARγ mRNA) | iFAT | 1.0 | 0.968 | 0.475 * | 0.242 * |
| PPARγ mRNA | oFAT | 1.0 | 3.260 | 0.519 ** | 0.292 ** |
| ASC-1 mRNA | iFAT | 1.0 | 1.100 | 0.193 ** | 0.366 * |
| ASC-1 mRNA | oFAT | 1.0 | 4.290 * | 0.330 | 0.433 * |
| AdipoR2 mRNA | Liver | 1.0 | 0.768 * | 0.718 ** | 0.427 ** |
| PPARα mRNA | Liver | 1.0 | 0.876 | 0.661 | 0.739 * |
| PPARα-Total.Protein | Liver | 1.0 | 0.600 | 0.782 | 1.43 ** |
| PPARα-phos-Protein | Liver | 1.0 | 2.630 * | 3.380 ** | 2.238 ** |
| PPARα.DNA.Binding | Liver | 1.0 | 0.583 * | 0.449 ** | 0.637** |
| HMGCR mRNA | Liver | 1.0 | 0.551* | 3.010 * | 1.030 # |
| ACCα.mRNA | Liver | 1.0 | 0.145 * | 0.747 | 0.168* |
| ACCα Total Protein | Liver | 1.0 | 0.987 | 1.923 * | 1.472 |
| ACCα.phos-Protein | Liver | 1.0 | 0.291 * | 0.34* | 0.681 |
| FFA | Serum | 1.0 | 0.962 | 0.683 | 1.656 # |
| FABP4 mRNA | Liver | 1.0 | 1.011 | 1.106 | 1.844 ** , ## |
| TNFα mRNA | iFAT | 1.0 | 1.897 | 5.820 ** | 8.032 * |
| TNFα mRNA | oFAT | 1.0 | 3.020 * | 2.54 * | 5.52 ** # |
| CD11b.mRNA | Liver | 1.0 | 1.197 | 2.401 ** | 3.787 ** |
| ALT Secreted | Serum | 1.0 | 1.556 ** | 1.733 * | 1.542 * |
| ALT mRNA | Liver | 1.0 | 1.855 * | 1.481* | 1.516 ** |
| IRS1 mRNA | iFAT | 1.0 | 1.074 | 0.542 ** | 0.403 ** |
| IRS1 mRNA | oFAT | 1.0 | 1.03 | 0.204 * | 0.208 * ## |
| Glut4 mRNA | iFAT | 1.0 | 0.815 | 0.546 * | 0.293 ** |
| Glut4 mRNA | oFAT | 1.0 | 0.956 | 0.239 * | 0.164 * # |
| IRS1 mRNA | Liver | 1.0 | 0.655 | 0.421 * | 0.369 ** |
| Glycogen | Liver | 1.0 | 0.830 | 0.787 | 0.822 |
| Insulin Receptor | Gastro | 1.0 | 0.778 | 0.575 * | 0.713 * |
| Glycogen | Gastro | 1.0 | 0.694 | 0.547 | 0.431 * |

\* $p < 0.05$ , \*\* $p < 0.01$ when compared to Sham-ConD; # $p < 0.05$ , $p < 0.01$ when compared to SCI-ConD.

